# Seizure dynamotype classification using non-invasive recordings

**DOI:** 10.1101/2023.04.02.535246

**Authors:** Miriam Guendelman, Rotem Vekslar, Oren Shriki

**Affiliations:** Department of Cognitive and Brain Sciences, Ben-Gurion University of the Negev, Beer-Sheva, Israel; Department of Computer Science, Ben-Gurion University of the Negev, Beer-Sheva, Israel

**Keywords:** EEG, Dynamotypes, Bifurcations, Blind source separation, Dynamical systems, Epilepsy, automated seizure classification

## Abstract

**Objective:** Recently, a seizure classification approach derived from complex systems and nonlinear dynamics has been suggested, termed the “taxonomy of seizure dynamotypes.” This framework is based on modeling the dynamical process of the transition in and out of a seizure. It has been examined in computational and animal models *in-vitro* and recently in human intracranial data. However, its applicability and value in surface EEG remain unclear. This study examined the applicability of dynamotype classification to seizure information extracted from surface EEG and tested how it relates to clinical factors.

**Methods:** Surface EEG recordings from 1,215 seizures were analyzed. We used an automated pre-processing pipeline, resulting in independent components (ICs) for each seizure. Subsequently, we visually identified ICs with clear seizure information and classified them based on the suggested taxonomy. To examine the possibility of automatic classification, we applied a random forest classifier combined with EEG features and evaluated its performance in identifying seizure-related ICs and classifying dynamical types. Lastly, we used a Bayesian estimator to examine the likelihood of the different dynamical types under various clinical conditions.

**Results:** We found an apparent onset and offset bifurcation in 49.5% and 40.3% of seizures, respectively. Bifurcation prevalence aligns with that previously reported using intracranial data and computational modeling. The automated classifiers, evaluated with a leave-one-patient-out paradigm, provided good performance. In addition, bifurcation prevalence differed between vigilance states and seizure classes.

**Significance:** We demonstrated a method to extract seizure information and classify dynamotypes in non-invasive recordings with a visual as well as an automated framework. Extending this classification to a larger scale and a broader population may provide further insights into seizure dynamics.

**Key Points Box:**

- We characterized the dynamical types of transitions at seizure onset and offset based on seizure information extracted from surface EEG.
- We classified the dynamical types (dynamotypes) in 49.5% and 40.3% of seizure onsets and offsets, respectively.
- The dynamotype distribution in surface EEG data aligns with previous findings from intracranial EEG and theoretical expectations.
- The likelihood of the dynamical type of a seizure exhibits differences across clinical seizure classes and vigilance states.
- Automated detection and classification of seizure bifurcations are possible using relevant features and pre-existing tools.

## Introduction

Epileptic seizures are classified based on clinical factors^1^. One limitation of this classification is that it does not relate to mechanisms of the ictogenic process. In view of the brain as a dynamical system, epilepsy can be considered as a condition with multiple stable states, having at least one interictal state and one synchronous oscillatory ictal state^2^. In such a system, a seizure’s start, evolution, and end are transitions between these states, called **bifurcations**^3^. Classifying seizures by how they dynamically start, and end may provide complementary information on the nature of different seizures^4^. While a classification method based on seizure onset morphology was previously suggested^5^, it does not connect seizure morphology to its underlying dynamical process. The “taxonomy of seizure dynamotypes” (TSD)^6^ is based on a minimal descriptive model, creating a classification framework that includes four possible onset and offset bifurcations, forming 16 different possible dynamotypes that could manifest in the electroencephalographic (EEG) signal^7,8^. The presence of all dynamotypes has been demonstrated in computational models and animal and human invasive recordings^7,9^. Saggio et al.^8^ first confirmed the existence of all bifurcation types in human data, showing that their empirical proportion is inversely related to their mathematical complexity, as expected theoretically.

Intracranial recordings provide a relatively clean signal from the epileptogenic zone (EZ), allowing identification of clear patterns. However, these recordings require an invasive procedure not performed in most patients with epilepsy. In contrast, surface EEG is non invasive and relatively low cost. Moreover, it is the most prevalent electrophysiological recording technique used in clinical diagnosis and follow-up of patients with epilepsy. It provides a complementary view to the invasive local recording, adding information on seizure propagation and evaluating the taxonomy of dynamotypes on a broader population. However, non-invasive recordings include multiple physiological and non-physiological sources, requiring their disentanglement and extraction of “clean” seizure-related information before applying dynamical classification.

This work examines one possible workflow to extract dynamotype information from surface EEG data. To this end, we used a combination of principal component analysis (PCA) and independent component analysis (ICA) to extract seizure information from the EEG signal. PCA has been applied to EEG signals to refine seizure classification performance^10^ and as a pre-processing step for source localization in ictal recordings^11^. The principal components (PCs) are linearly independent of each other and orthogonal^12^, which will not optimally reveal the seizure sources. To refine the sources contained in this signal, we applied Infomax ICA^13^.

In the context of EEG analysis, it is typically used both in pre-processing for source analysis and de-noising^14,15^. ICA was used in previous studies for seizure detection, classification, and identification of the seizure onset zone, with surface EEG^16–20^ and magnetoencephalography (MEG)^21–23^ achieving good performance. Barborica et al. found a strong correlation between cortical seizure sources in intracranial EEG and the estimated sources evaluated from surface EEG using ICA.^16^ Recently developed tools may support automation of the classification process. We used these algorithms to score and classify the independent components (ICs) into brain and non-brain sources^24,25^, identifying physiological artifacts such as muscle-, heart-, and eye-related artifacts and external artifacts such as line and channel noise.

An abrupt system change, such as a bifurcation, can also be described as a change point^26^. Existing tools for automated change point detection^26^ can provide more accurate temporal identification of the transitions into and out of a seizure. Moreover, employing features used for seizure detection^27^ together with interpretable machine learning^27^ allowed us to create and evaluate an automated classification process.

### Objectives of current work

Our primary objective was to evaluate our ability to classify seizures visually using TSD in non-invasive EEG recordings. Second, we examined the use of existing tools in assessing visual labeling quality and developing an automated seizure information identification and classification system. Third, we evaluated the dynamotype proportion in our data and compared it to previous findings and the expected dynamotype ratio. Lastly, we aimed to quantify and compare the distribution of different bifurcations in sub-groups of patients (by age, gender, etiology, or localization) and seizure characteristics (by clinical classification or vigilance states).

## Methods

### Data

We used the EPILEPSIAE database^28^, containing annotated long-term surface EEG recordings from 158 patients with intractable epilepsy undergoing pre-surgical evaluation at three medical centers. Two experienced EEG reviewers annotated the data. Patient metadata included: gender, age, etiology, and EZ when identified. Seizure annotations included onset and offset times, vigilance state, and seizure classification. Seizure classification in the database was according to the 1981 classification scheme^29^, including simple partial, complex partial, and secondarily generalized, respectively, matching focal aware (FAS), focal with impaired awareness (FIAS), and focal-to-bilateral tonic-clonic (FBTCS)^1^. The uncategorized (UC) seizure does not directly translate into the new classification; hence, it remained as is [see Ihle et al. (2012)^28^]. A total of 1,215 seizures were analyzed. The data consisted of 19 to 37 electrodes using the 10-20 placement system, with a 256–1024 Hz sampling rate. We excluded one patient with corrupt recordings during seizure times.

### Pre-processing

The analyzed data included a margin of 30% of the seizure length at the beginning and end of each seizure. For each seizure, we randomly selected a control segment of similar size at least five minutes away from a seizure in the same recording. If there were insufficient data for a control segment, we omitted the seizure (n=38). Finally, we used 1,177 seizures for subsequent analysis. We built an automated pipeline to clean and decompose the EEG signal (both seizure and control) using EEGLAB functions and a costume code. The pipeline included: downsampling to 256Hz, highpass filtering at 1Hz, artifact subspace reconstruction^30^ with bad channel rejection and interpolation, re-referencing to the mean, and lowpass filtering at 40Hz. We then applied PCA, keeping the first eight PCs, followed by ICA, resulting in eight ICs for each seizure.

### Labeling dynamotypes

Two independent reviewers evaluated each seizure component (n=9416) for the onset and offset bifurcations (Figure 1A). Using our in-house labeling application, the reviewer annotated bifurcations with clear morphology (Figure S1). We used the visual classification flow described by Saggio et al.^8^. Most surface EEG recordings do not include DC (zero frequency) component recording, limiting the ability to distinguish between several bifurcation types. Specifically, it is impossible to distinguish Saddle Node (SN) and Subcritical Hopf (SubH) bifurcations at onset, nor the Saddle Homoclinic (SH) and Saddle Node on Invariant Circle (SNIC) at offset. In this work, we regard them jointly, as was previously done^8^. The onset labels included: SupH, SNIC, SN/SubH, and unclear, and the offset labels included: SupH, SNIC/SH, FLC, and unclear. In Figure 1, panels B and C provide examples of each onset and offset bifurcation classified in our empirical data. For additional information on the labeling protocol, interface, and examples, see Figures S2, S3, and Appendix 1.

**Figure 1:**
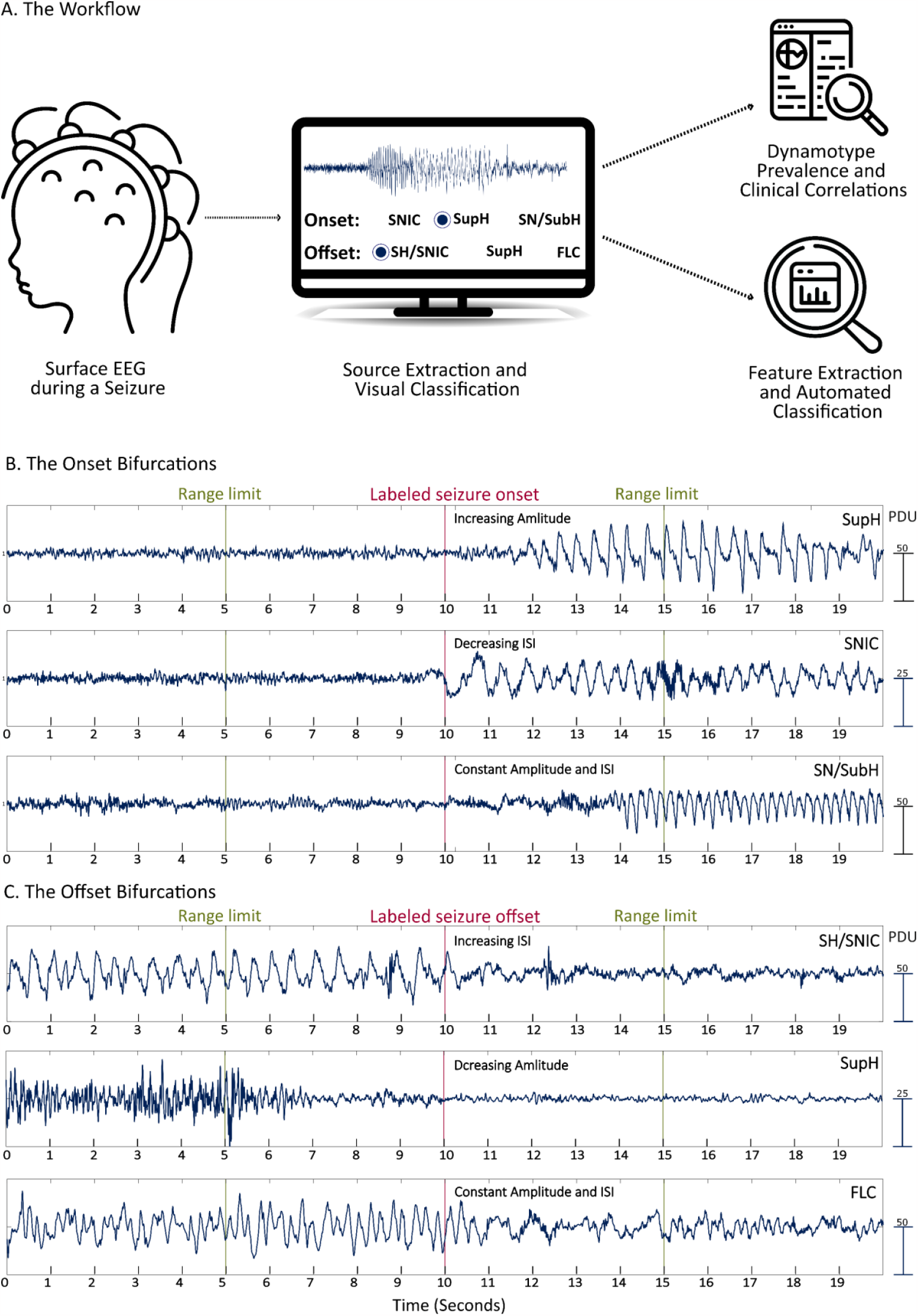
Workflow and bifurcation examples. **(A)** An illustration of the workflow. Surface EEG data were automatically pre-processed, and the independent components were presented to the reviewer. The reviewer then labeled each component with a bifurcation or as “not clear” if no clear bifurcation could be identified in the time range of 10 seconds around the original seizure annotation. Subsequently, the labels were analyzed with two aims: 1) the relationship with clinical factors and 2) the examination of features for automated classification. **(B)** Examples of the three onset bifurcations: First, we can see SupH, accompanied by the amplitude increase, and SNIC, characterized by larger inter-spike intervals (ISIs) close to the bifurcation. Lastly, SN and SubH, differing only by a DC shift, are indistinguishable in our recordings. The amplitude and ISIs remain constant after seizure onset. The scale of the components is affected by matrix pre-whitening before the PCA and ICA procedures; therefore, the units are presented as procedure-determined units (PDUs). **(C)** Examples of the three offset bifurcations: Similar to the onset bifurcations, SupH exhibits an amplitude increase, whereas SNIC and SH are characterized by an increase in the ISIs towards the seizure’s end. However, the lack of DC in our recordings makes them not distinguishable. Lastly, FLC is displayed with a constant amplitude and frequency.

### Label validation and quantitative analysis

#### I. Inter-rater agreement evaluation

We evaluated for inter-rater agreement using percent agreement, Cohen’s Kappa coefficient, and confidence intervals^31^. This evaluation was made separately for the onset and offset labels under two conditions: selecting seizure information and agreement in mutually chosen components.

#### II. Establishing that selected components are of brain origin

We scored all components using Iclabel^24^ and MARA^25,32^. The score was calculated globally for each component and locally around the onset and offset bifurcations. The 1-MARA score and Iclabel brain score indicate the probability of a component being of brain origin. We evaluated the median score in three groups: seizure ICs with a clear bifurcation (onset or offset), unselected seizure ICs, and control ICs.

#### III. Change point detection for bifurcation time detection

We evaluated the ten seconds around the dataset annotations (onset and offset) for the seizure and corresponding times in the control data. We used MATLAB’s built-in function, “***ischange***,” a variance-based change point detection, and limited detection to a ***single change point***, analyzing the proportion of detected change points in the three groups. In cases with a clear change point, we used it as an anchor for the subsequent feature extraction. This anchor strongly impacted feature quality, especially for estimating amplitude and inter-spike interval (ISI) scaling.

#### IV. Component selection

In some cases, we labeled more than one IC as having a clear bifurcation. In most cases, the components for each seizure included the same dynamical class. Nevertheless, 40/583 seizures at onset and 53/474 at offset had ICs labeled with more than one bifurcation. In these cases, we determined the seizure bifurcation by a simple decision flow, illustrated in Figure S4. We designed this flow to select a component with a decent brain score (> 0.6) and the earliest change point detected in the seizure.

### Bifurcation and dynamotype distribution

We first evaluated the overall bifurcation detectability and dynamotype proportion on all seizure data. Second, we summarized the number of different bifurcations observed in individual patients in the context of the overall bifurcations observed. Third, we wanted to estimate the bifurcation probabilities in the context of various clinical factors. To assess these, we used a Bayesian multilevel multinomial logistic model (brms package in R^33^) to account for repeated measures within patients and multiple clinical factors. The model included age, gender, etiology, EZ localization, clinical classification, seizure length, and vigilance state. For the onset, we used all three bifurcation labels. Because we had only three SupH cases for the offset labels, we focused on the distinction between cases with or without ISI slowing. We then evaluated the difference in each bifurcation’s posterior distribution, given the different clinical factors. We used the probability of direction (pd) and maximum a-posteriori-based p-value (MAP-P) to evaluate distribution differences^34^. Full region of practical equivalence (ROPE) was defined as an effect size smaller than a 20% relative change and used to estimate the effect magnitude. For more details on the model, priors, and metric considerations, see Appendix 3.

### Feature extraction and automated classification

#### I. Feature extraction

We used brain score values and the presence of a change point as features. Using the change point time as the starting time, we extracted statistical, spectral, autocorrelation, entropy-based, and morphological features quantifying amplitude and ISI change. We omitted components with unreliable features from the analysis. For more details on this process, see Appendix 4.

#### II. Automated classification

We separated this task into two steps: first, identifying seizure-related components, and second, distinguishing seizure components with or without ISI slowing. For each task, we used the complete set of features. We used a random forest classifier^35^ with two cardinal parameter settings: tree depth of three to avoid overfitting and class weights to the inverse of the class proportion during training to counter the effect of class imbalance. We evaluated the models using a leave-one-patient-out cross-validation paradigm. Finally, we performed a SHAP^27^ analysis to gain insight into the contribution of the features to each classifier. (For more details, see Appendix 5.)

## Results

The present study aimed to investigate the dynamical type of seizure bifurcations in 1,177 seizures collected from 158 focal epilepsy patients during pre-surgical evaluation. We processed multi-channel ictal EEG data using PCA and ICA to produce independent source signals (see Methods). The results were visually classified, and we examined the overall proportion of seizure bifurcations and their interaction with clinical factors. Additionally, we propose an automated classification pipeline for more extensive population scalability.

### Label Validation

#### I. Inter-rater agreement evaluation and overall detectability

Both reviewers evaluated 1,177 seizures and a total of 9,416 components. Of these components, each reviewer selected and labeled only those with apparent seizure information with a clear onset or offset bifurcation. The overall percent agreement for defining apparent onset and offset bifurcations was 90.04%, 90.1%, Cohen’s kappa 0.657 [CI 0.637–0.678], and 0.641 [CI 0.618–0.663], respectively. From all the components, both reviewers selected 1,132 and 948. The percent agreement on bifurcation type within the selected components was 97.44% for onset and 93.25% for offset, Cohen’s kappa 0.88 [CI 0.84–0.92], 0.84 [CI 0.80–0.88], respectively, which is considered substantial^36–38^. We used only mutually selected components with an agreement for further analysis (1,103 for onset, 884 for offset), and defined components without agreement as unclear. For additional information on chosen components, see Appendix 2.

At the seizure level, we found at least one onset and offset bifurcation in 49.5% (n=583) and 40.3% (n = 474) of the seizures. Of all seizures, 362 had both an onset and offset label, using these for the dynamotype analysis (221 had only a clear start, 112 had only a clear end, and 482 had neither). Bifurcation detectability was better for more posterior EZ and worse for frontal (Figure S5A). Interestingly, seizures with a more extensive clinical manifestation were related to a more significant number of labeled bifurcations (FBTCS > FIAS > UC > FAS, Figure S5B).

#### II. Change point detection

We analyzed the proportion of identified change points in the three groups around seizure annotation times or matching times in the control segments. The proportion of segments with a detected change point was the lowest for control recordings (51.5%, 51.1%), intermediate for the non-selected seizure components (63.5%, 61.9%), and the highest for the selected seizure components (77.9%, 84.3%).

#### III. Brain score

Globally, MARA and Iclabel brain scores were highly correlated (r=0.66, p<0.001 in the control data, and r=0.6, p<0.001 in the seizure data). We calculated mean scores for each component and referred to it as the “score.” The median scores are 0.62 [IQR: 0.30–0.94] for all seizure data and 0.81 [IQR: 0.42–0.984] for the control data, which we expected considering the relatively large amount of noise expected during a seizure. The components selected by the raters had scores (median: 0.92 [IQR: 0.75–0.98]) that were higher compared to the non-selected ones (0.53 [IQR: 0.243–0.91]), supporting the selection of brain information in our dynamotype classification. We note that components with a high brain score can include non-ictal or ictal activity without a clear bifurcation morphology in the defined time range (Figures 2SB–E). See Appendix 2 for an additional validation analysis.

### Dynamotype prevalence

#### I. Onset and offset bifurcation proportion

First, we individually analyzed the onset and offset bifurcation proportion (Figure 2A). Importantly, our findings were consistent with previous ones regarding the overall rank^8^. The dominant onset type was SN/SubH (88.7%), the second was SupH (6.7%), followed by SNIC (4.7%). For the offset, FLC (70.7%) was most prevalent, followed by SH/SNIC (28.7%), and then by SupH (0.6%), which was rare in our data. On average, patients with more seizures had more bifurcation types, suggesting that given a more significant number of seizures, it is likely to observe a broader range of bifurcation types in each patient (Figure 2B).

**Figure 2:**
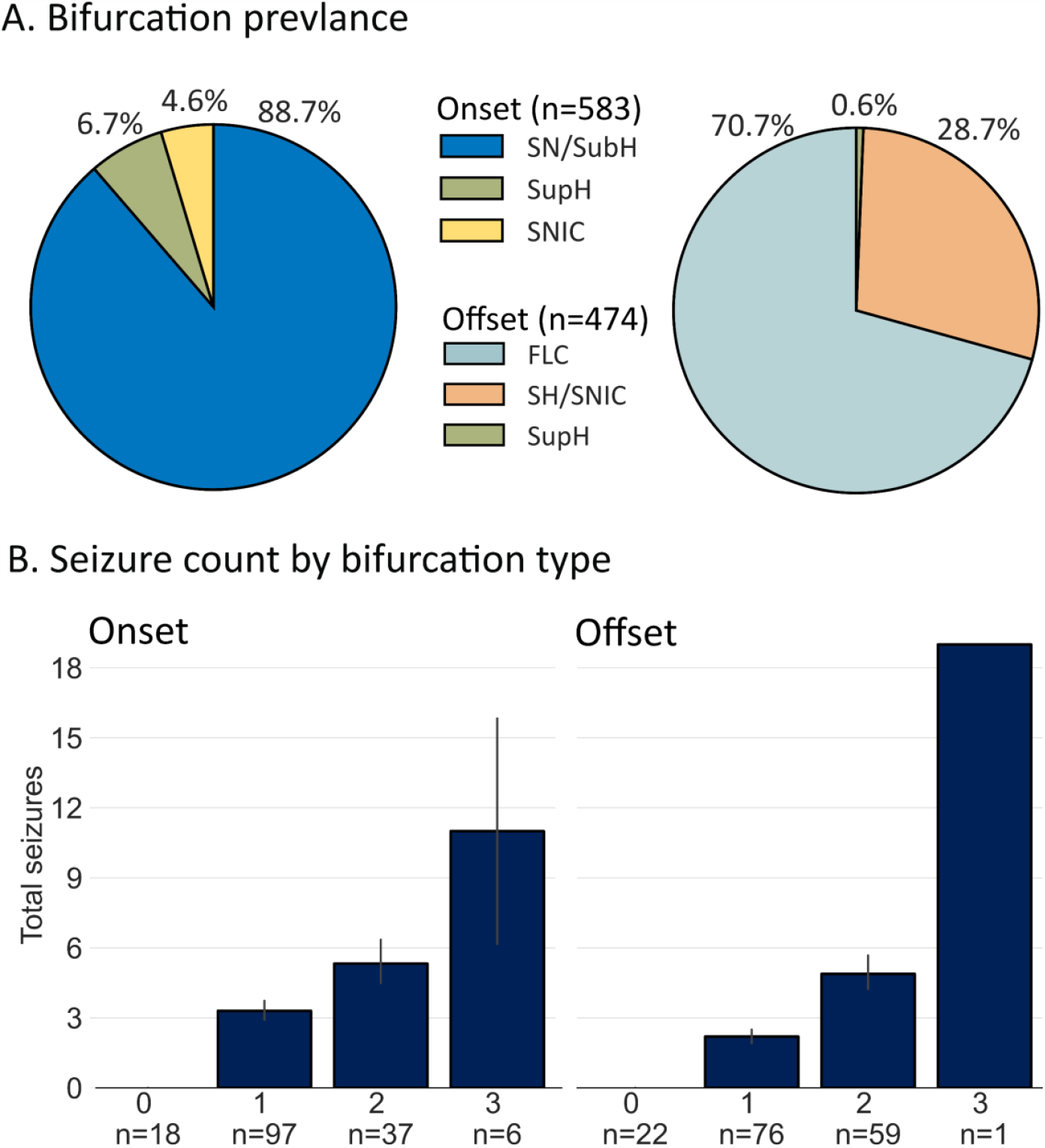
Dynamotype distribution and within-patient occurrence. **(A)** The bifurcation proportion found in our data using all seizures labeled with onset or offset bifurcations. The most prevalent onset bifurcation is SN/SubH, and the most prevalent offset bifurcation is FLC. **(B)** The mean number of classified onset or offset bifurcations in a patient is a function of the number of bifurcations found within that patient. We can see that in patients with more seizures, we could find more bifurcation types, suggesting that, given additional data, it is likely to find more than one onset and offset bifurcation in each patient. This was demonstrated for seizure offset regarding ISI slowing in previous work using long-term recordings^8,47^.

#### II. Dynamotype prevalence

Dynamotype complexity is determined mainly by the minimal co-dimension (number of parameters that need to be tuned to obtain a dynamotype) and number of bifurcations required to produce such a dynamotype^7^. Table S3 summarizes the dynamotype count found in our data, ranked according to the estimated complexity described by Saggio et al. ^7^. Consistent with theoretical expectations, the most prevalent dynamotypes were the least complex (SN/SubH – FLC, n=241; SN/SubH – SH/SNIC, n=83). More complex dynamotypes were scarce (0–14), requiring a larger sample for estimating the relationship with theoretical complexity.

### Relationship with clinical factors

In the clinical factors analysis, we report results for groups with at least 20 seizure examples. We consider pd > 0.99 or p-MAP < 0.01 as a statistically significant difference. Considerable effect magnitude was determined by ROPE < 0.03 (and any effect practically null by ROPE > 0.97). For extended statistical testing methods, see ***Appendix 3***. Regarding the vigilance state, the overall bifurcation proportion was similar within the NREM stages, with a small proportion of seizures not beginning with SN/SubH (Figure 3A). The wake and REM stages had larger SNIC and SupH onset proportions than the NREM stages. We found a substantial difference when comparing the SNIC proportion between awake and deep sleep (pd = 0.9975, p-MAP = 0.0002, ROPE = 0.0088).In the clinical classification, FIAS and UC seizures had a higher proportion of SupH than FBTCS and FAS. The ratio of SupH in UC (10.9%) and FIAS (6.2%) was more prominent than in the FBTCS (0%) and FAS (1.8%) groups (Figure 3B). This difference was significant for FBTCS compared to FIAS or UC seizures (pd > 0.99, p-MAP < 0.001, ROPE = 0.01 for both) (Figure 3B). Our results did not indicate a substantial difference in the ISI slowing proportion in the different clinical conditions. Other trends appearing in the data did not reach our significance thresholds; for detailed information, see ***Supporting Tables 1 & 2***.

**Figure 3:**
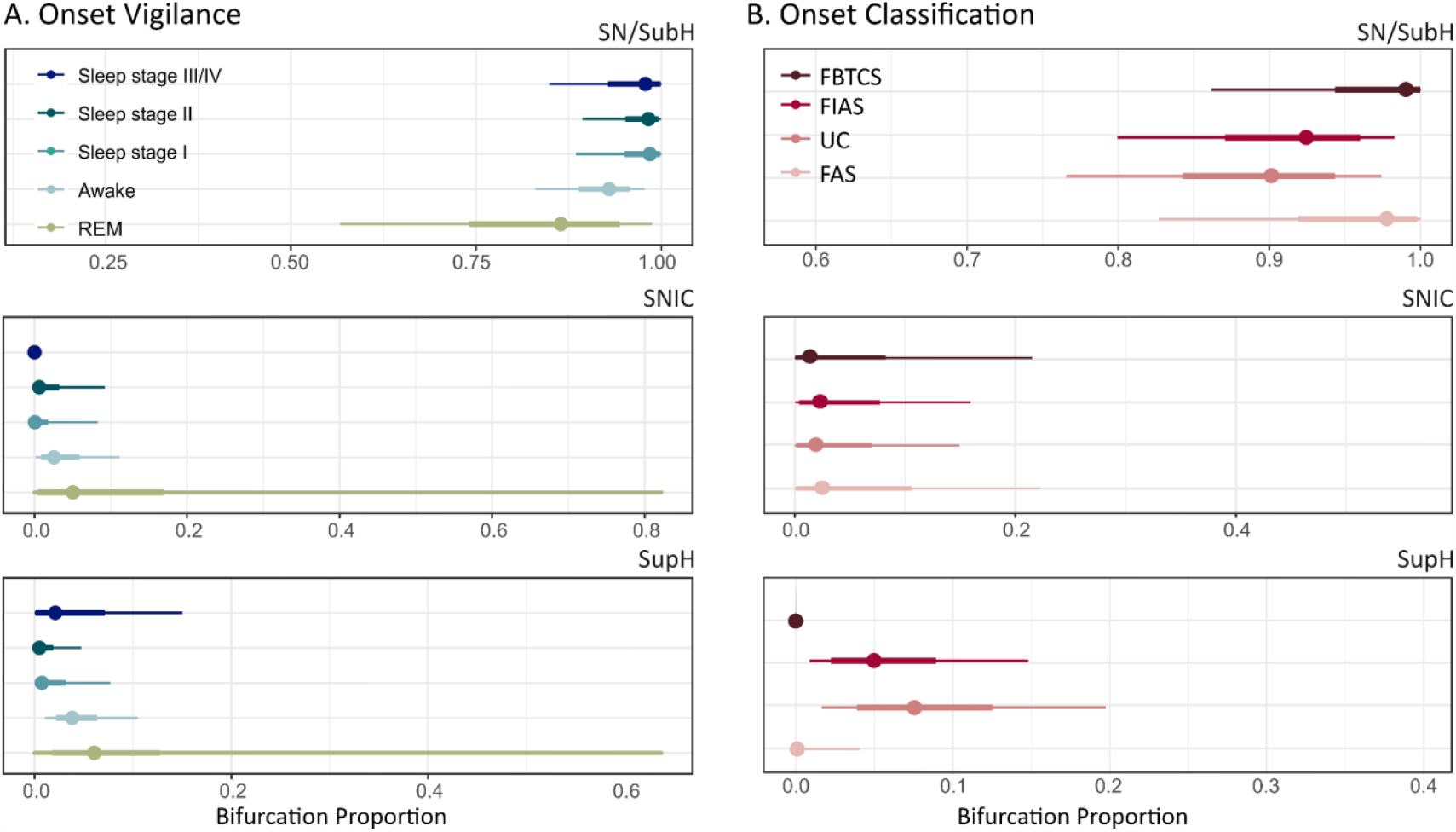
The different dynamical distributions in clinical factors. **(A)** The distribution of all three onset bifurcations across the vigilance state shows nearly only SN/SubH bifurcations in NREM sleep. SNIC and SupH are slightly more prevalent in the awake and REM stages. **(B)** The estimated distribution of all three onset bifurcations as a function of seizure classification. We can see that SupH bifurcations are more prevalent in FIAS and UC seizures.

### Automated classification of components and bifurcations

Automating the classification is crucial for the scalability and consistency of the dynamotype analysis. Of the 9,416 components analyzed, we visually labeled a clear transition in 1,103 onsets and 884 offset components. We omitted components with unsuccessful feature extraction (685 onsets and 742 offsets); of these, only a small number had a clear label (4 onsets, 19 offsets); see Appendix 5 for additional details. For each classification task, we used a random forest classifier^39^. We evaluated performance using a leave-one-subject-out paradigm, using balanced accuracy (BA)^40^ and the area under the receiver operating curve (AUC-ROC).

#### I. Detecting a seizure component

Altogether, 8,731 onset and 8,674 offset components were included; 1,099 onsets and 865 offsets had a clear bifurcation. The median BA was 73.3% [IQR: 66.7-81.6%], n=158, and AUC-ROC was 0.83 [IQR: 0.76-0.90], n=148. Figure 4, left panel, presents the top ten features affecting the model’s decision. Six top features were related to scores based on the MARA and IClabel classifiers (score global, brain global, score local, brain local, max score, and other class global). Low values in these features tipped the classifier’s decision to the “unclear” label, yet positive values show a weak relationship with the “clear” label. Additionally, three entropy measures used for seizure detection^41^ were ranked as the top features affecting this task. Lastly, a higher mean peak amplitude supported the decision towards seizure information.

**Figure 4:**
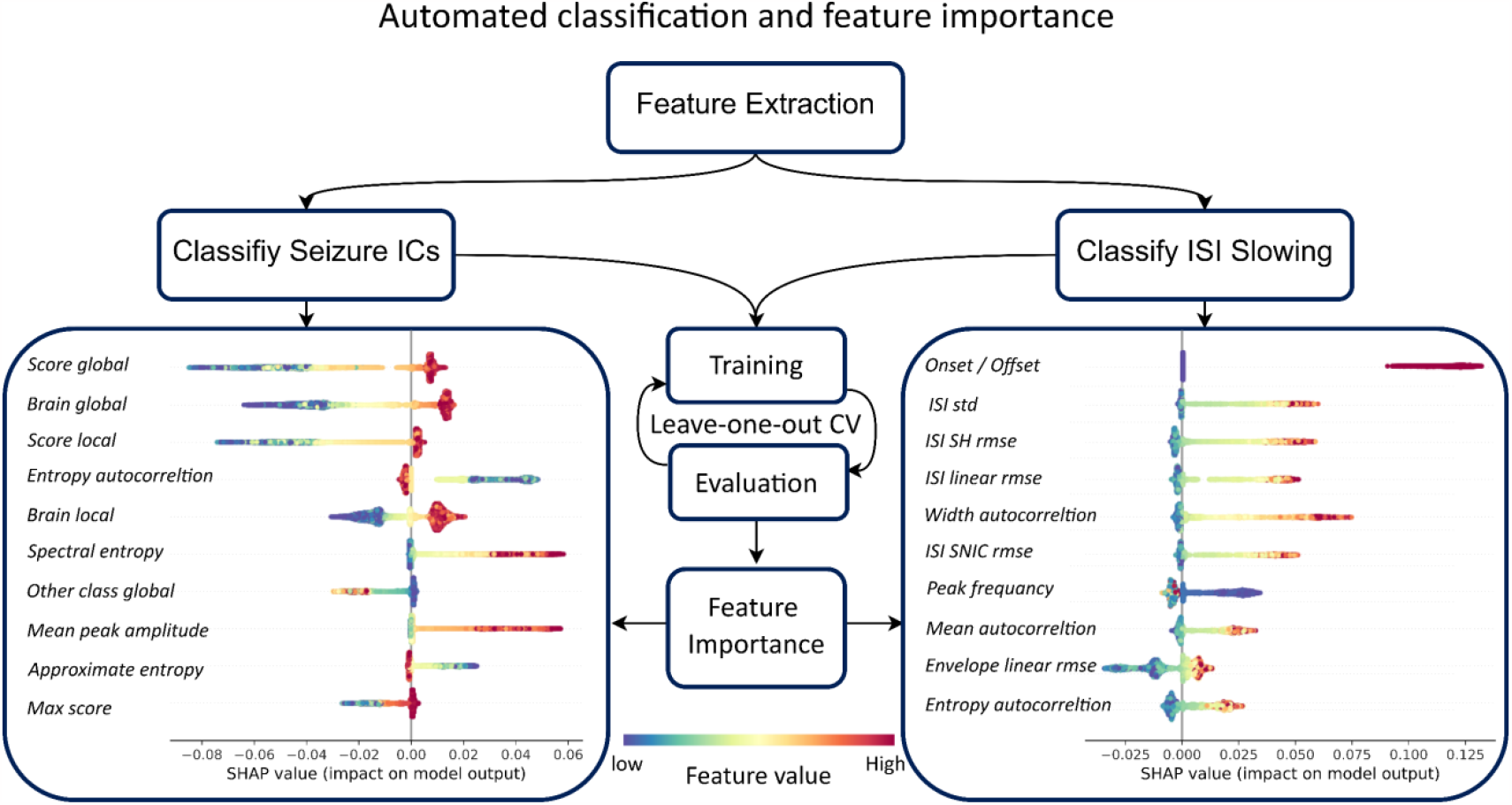
Feature analysis and automated classification. This figure demonstrates the classification workflow and feature importance. The first features were extracted for both onset and offset bifurcation, as specified in Appendix 3. All features were employed for two classification tasks: Identifying the seizure-independent components (ICs) and classifying inter-spike interval (ISI) slowing towards the bifurcation. Each model was trained and evaluated using leave-one-patient-out cross-validation (CV), and the balanced accuracy distribution was calculated. Finally, we trained the model on all the data and performed SHAP analysis^27^. In the figure below, we can see the top ten features contributing to each task.

#### II. Classifying ISI modulation

In this stage, 1,099 onset and 865 offset components were used: 52 onsets and 234 offsets had ISI slowing toward the bifurcation. The median BA was 80% [IQR: 64.96-95.25%], n=148, and AUC-ROC was 0.91 [IQR: 0.8-1], n=86. Figure 4, right panel, presents the top ten features affecting the model’s decision. The chief feature was the onset/offset indicator, as ISI slowing was more probable for the offsets in our data. Features quantifying ISI changes and autocorrelation features had the highest impact on this classifier decision. The strongest within these was the width of the autocorrelation function, previously related to critical slowing at larger scales^42^.

## Discussion

Recent studies using dynamical models^9^ combined structural, anatomical, and functional data to create a digital simulation of the brain network^43^ replicating seizure patterns^44,45^ in patients undergoing pre-surgical evaluation and improving surgical outcomes^46^. Similar models have defined the “taxonomy of seizure dynamotypes.” This taxonomy has helped characterize changes in seizure dynamics along the epileptogenic process in animal data^47^. In human data, this classification has been explored in invasive electrographic recordings in focal drug-resistant patients^8^. Extending this classification to non-invasive EEG recordings allows us to examine seizures dynamically on a broader range of epilepsy syndromes, stages, and provocations.

The challenges in such an analysis go beyond the noise in the surface EEG signal. The non-invasive signal records a more extensive population of neurons such that a single electrode may record both the seizure onset and propagation zone. Additionally, the neuronal signal is transmitted through the CSF, dura matter, skull, and skin, subjecting it to volume conduction and smearing^48,49^. Surface EEG has limitations, but the demonstration that similar seizure dynamics are observed at different scales and with a similar prevalence supports the view of scale-free system behavior. The present study explored this classification in a large surface EEG dataset^28^ with high-quality seizure annotation. Using our approach, we identified a clear dynamical type in 49.5% (n=583; onset) and 40.3% (n = 474; offset) of seizures.

In general, the overall bifurcation ranking was aligned with previous findings in invasive recordings^8^. Interestingly, the proportion of dynamotypes ending in FLC was higher than expected (70.7%, compared to the previous finding of 53–54%^8^). The FLC might be favored by the need for terminating oscillations in large-scale neuronal networks^50^.

From a dynamical point of view, we consider the ictal and non-ictal states as dynamically different. Beyond these states, the awake and sleep stages from which seizures arise are also considered different dynamical states^51^, potentially affecting seizure dynamics. In Figure 4A, we see that seizures occurring in NREM sleep had nearly only SN/SubH bifurcations. In contrast, the awake and REM states had more variability in onset morphology with a more significant proportion of both SNIC and SupH bifurcations. Only the difference in the SNIC proportion between awake and NREM3 reached our statistical criteria, but this could be due to a low number of examples in the REM stage. Moreover, the staging in these data was based on the sleep stage 30 seconds before the seizures. An analysis in a broader temporal context may reveal a stronger relationship with vigilance states.

We found differences in SupH onset when comparing seizure classes. In surface EEG, when evaluating the SupH morphology, the amplitude increase may correspond to propagation or conduction. It may not reflect the pure dynamics of the neuronal population in the onset zone. In some cases, we could identify amplitude increases that seemed to be related to the propagation and not the onset dynamical type (Figure 1SB–C). Although we tried to separate such cases, propagation may have been misclassified as SupH. Interestingly, we did not observe SupH in seizures classified as FBTCS, where propagation is mainly expected. Future work using simultaneous intracranial and surface recordings could help provide insight into the expression of SupH in surface EEG.

Regarding limitations, our pipeline was significantly less sensitive to more localized seizures (FAS) and frontal seizures, typically less prominent in surface EEG data. An additional theoretical limitation is that our pipeline assumes seizure information comprises high-amplitude synchronized activity. Hence, seizure onset comprised of low voltage fast activity will potentially be missed^5^. These seizures may be better identified using different processing pipelines with different pre-processing approaches. Yet, in many cases, we observed seizure components starting with low-amplitude fast activity, evolving to high-amplitude low-frequency activity in the same component, which resembles the evolution of a seizure in the epileptogenic zone as previously described^5^.

The visual classification process is labor intensive and error prone. For more consistent results and label validation, we used two independent labelers and several validations specified in the “label validation section.” Development of automated tools that can identify the seizure information and classify its dynamical type will allow us to extend this analysis to larger cohorts and a wider population. In this work, we implemented a baseline classification algorithm to examine the value of features for these tasks, reaching good performance in both. We demonstrated that the top features contributing to identifying seizure information included brain scores and entropy measures typically used for seizure detection^52^. The ISI slowing classification relied on ISI morphological and autocorrelation features, previously linked to critical slowing down^42^. Combining a more extensive set of labeled data, a more specific change-point approach^53^, and a larger feature set^52^, and examining different machine-learning approaches may result in more sensitive and specific classifiers. Current and previous work has focused on seizure onset and offset bifurcations. Automated detection tools, particularly change point analysis throughout a seizure and automatic classification tools, may allow us to quantify the dynamics throughout an entire seizure^53,54^.

## Conclusions

We demonstrated that applying the TSD to surface EEG is possible, albeit with certain limitations. The overall dynamotype distribution aligns with previous findings and theoretical expectations. Additionally, we found that bifurcation proportions may be affected by vigilance and seizure classes. Using our visual labels, we created two automated classifiers for identifying seizure information and classifying ISI slowing, showing promising cross-subject performance. We evaluated the feature contribution for each task, suggesting the use of feature subsets in further development. Moreover, such automated classifiers can extend this analysis to a more extensive database, reducing the labeling workload as an initial screening step.

## Acknowledgments

This research was supported by grants 504/17 and 794/22 from the Israel Science Foundation (https://www.isf.org.il).

We want to acknowledge Matan Ben-Shahar for his statistical consultation, and Carmel Salomonski for her assistance in building the graphical user interface and conducting the initial analysis. Their contributions greatly enhanced the quality of our study. We are grateful for their expertise and support.

## Contributions

MG, RV, and OS conceived of the presented idea, MG and RV created the visual interface and performed the visual analysis, MG performed the statistical and automated classification analysis, and OS supervised the findings of this work. All authors discussed the results and contributed to the final manuscript.

## Disclaimer

The authors declare no conflicts of interest.

## Data availability statement

This worked used the data from the EPILEPSIEA database described in Ihle et.al. 2012^28^. All the project code base used to is available in the project <u>GitHub repository</u>. Additional data that support the findings, e.g. the component time series and visual labels of this study are available from the corresponding author upon reasonable request.

