## Supporting information for "Seizure dynamotype classification using non-invasive recordings"

### Appendix 1: The labeling interface, protocol, and examples

#### Labeling protocol

The EPILEPSIAE dataset[1] used for this project includes seizure labels for onset and offset times, as described in the main text. We used the reference onset and offset times and allowed a five-second range around the labeled time to identify a bifurcation. We then used this ten-second range to visually analyze each independent component's onset and offset. We first identified if there was an apparent bifurcation at the given time range as a clear onset or offset of the seizure pattern. We evaluated the clear bifurcations for amplitude and inter-spike interval (ISI) changes for five seconds from the identified bifurcation, as described in Saggio 2020[2]. Following a training session, two independent reviewers labeled all the resulting seizure components (n = 9416). After the first labeling iteration, in a discussion between reviewers, we concluded that the labeling process had a learning curve, especially the distinction between data originating in the brain or from other physiological sources. Labeling errors are less likely to have happened in components that were labeled as having a clear bifurcation by both reviewers (not necessarily labeled the same) or marked as unclear by both reviewers. We revised the components with no agreement, during which each reviewer relabeled these components **independently.** At this stage, the reviewers were blinded by the previous labels, and the new labels replaced the old ones in these seizures.

#### Labeling interface

Four panels were presented to the reviewer for each seizure using a graphical user interface (GUI). The panels include the EEG and the component time series, the component characteristics, a temporal and topographic activation map, and the power spectrum of the component during the entire seizure and a labeling user interface to store the labeling results using costume code combined with EEGLAB[3] visualization functions (Figure S1).

#### Labeling examples

To revise labels in the second step, we developed a more efficient complementary interface focusing only on the time series' morphology at relevant signal time points. This framework focuses separately on individual components and highlights the analysis range of either the onset or offset in seizures. With this interface, we could revise the labels more efficiently in the second step described above. For decision examples, see Figure S2 for seizure onset and Figure S3 for offset.

### Appendix 2: Validation of component selection

#### I. Inter-rater evaluation

Tables S1 and S2 are the confusion matrices of the components selected by both reviewers for onset and offset. After the agreement evaluation, we marked the label as unclear if both reviewers selected a component and did not agree on the bifurcation (not on the diagonal of the confusion matrix). Only seizure components independently selected by both reviewers with **agreement on the bifurcation** class were included in the subsequent analysis as clear bifurcations.

#### Score analysis

The independent components were sorted by the explained variability of the signal they account for, so the first component includes the largest proportion of explained variability. In addition to the general analysis, we were interested in examining this difference in the context of the component rank. In Figure S4A, we see the decrease in score difference in the low-ranking (high number) components. In addition, the proportion of components with a clear bifurcation was reduced in low-ranking components, Figure S4B.

### Appendix 3: Bayesian multilevel modeling and inference

Since multiple bifurcation types can occur in a patient, we should consider all seizure types found in each patient and their prevalence. Additionally, in this analysis, we should account for repeated measures and estimate the proportion in the context of all clinical parameters to avoid biases in the data. Thus, multilevel multinomial modeling is essential for proper statistical evaluation of the effect of these factors. Considering the large number of and the probabilistic nature of bifurcation occurrence, we constructed a multinomial multilevel model using the "Bayesian Regression Models using Stan" (brms) package in R[4]. We used brms default priors, which put flat priors on all fixed effects. The model distribution parameters were estimated using Monte-Carlo-Markov-Chain (MCMC); for each model, we used six chains with 7,000 iterations each; the first 2,000 iterations were considered a warm-up and not included in the final posterior estimation. This approach allowed us to estimate the posterior distributions of each bifurcation given our data in different clinical conditions.

We included clinical factors potentially affecting brain and seizure dynamics in our analysis. We modeled the data using age, gender, side, lobe, and etiology as patient-level effects. To account for within-subject repeated measures, we set the patient as a random effect for vigilance, seizure length, and classification. We used all three bifurcation categories, SN/SubH, SNIC, and SupH, and fit a categorical logistic regression model for the onset. For the offset, we had only three seizures with a SupH offset. Thus, we created a binomial model distinguishing bifurcation with (SH/SNIC) or without ISI slowing (FLC, SupH).

The Bayesian modeling approach allowed us to compare the posterior distributions explicitly and provide an intuitive understanding of the effective direction and size when doing such comparison[5]. In this work, we analyzed the ratio of two given posterior distributions to quantify the relative change in the probability of each class. Here, for the definition of equivalence, we should notice the proportion of the minority class 89%, 6.5%, 4.5% (for onset) and 72.1%, 27.9% (for offset); in each analysis, we defined practical equivalence as a change smaller than 20% relative to each class (that is, any ratio between [1/1.2=0.833 and 1.2]). Tables S3 and S4 summarize the results for all comparisons. In Bayesian statistics, we use specific measures when evaluating the difference between two distributions. In our analysis, we used the following:

**The probability of direction (pd)**—mathematically defined as the proportion of the posterior distribution at the median's side, dividing this distribution at the point of the null hypothesis; it typically ranges between 0 and 1. We consider pd > 0.99 as a significant directional effect[5].

The **maximum a posteriori (MAP)-based p-value**—relates to a parameter's odds against the null hypothesis. It is mathematically defined as the density value at the null hypothesis divided by the density value at the maximum a posteriori; here, we consider P-MAP values below 0.01 as significant.

To evaluate the difference between two conditions in a factor, for example, cortical dysplasia and temporal sclerosis in the etiology factor, we first calculated the posterior distribution of each bifurcation's expected probability for each condition marginalized across all other factors. To evaluate the difference between the distributions, we calculated a new posterior distribution, the distribution of ratios between the conditions, and evaluated the pd and p-MAP for the resulting distribution.

We used the full region of practical equivalence **(ROPE)** to evaluate the effect magnitude. This approach defines the null hypothesis regarding a range of values considered negligible or too small to be of practical relevance as the ROPE. Then we estimated the posterior probability of the estimated effect within the ROPE. ROPE probabilities > 0.97 indicate strong support for the effect being practically negligible, while a ROPE < 0.03 indicates a significantly meaningful effect size[5].

### Appendix 4: The extracted features

We used the change point found in the signal as the onset time. In cases where the change point was not found, the dataset annotation time was used as the start point. We flipped the offset signal over time for consistency with the onset signal. We standardized the ten seconds around the time of interest for both onset and offset to achieve a more stable feature range. We extracted all features using a MATLAB costume code.

***Morphological features:*** Visual bifurcation classification is based on typical morphological features, mainly amplitude modulation and Intra spike interval (ISI) modulation. For these features, the bifurcation timing critically affected the feature quality. The most coherent results were achieved with two seconds for amplitude modulation and five seconds for ISI modulation. We used the individual peak amplitude and the signal's envelope root mean square (RMS) estimation for amplitude modulation. We calculated the ISIs as the time gaps between the identified peaks. We set a typical morphological curve for each value as previously described[2,6,7]. We then fitted and evaluated using the fitting parameters, root mean square error (RMSE), and adjusted R squared for each curve fit. ***Statistical features:*** We calculated the mean, median, standard deviation, and skewness of the signal over five seconds from the seizure onset point.

Additionally, for the ISI, the mean distance and STD were calculated. ***Spectral features:*** The signal's power spectrum was calculated between 1–40Hz, and a linear fit between the logarithm of the power spectrum as a function of frequency was estimated. The intercept, slope, RMSE, and adjusted R squared were estimated. The furthest positive point from the linear fit was identified as the frequency peak, expected to be related to the dominant seizure oscillatory frequency. ***Autocorrelation and entropy-based features:*** Spectral entropy and approximate entropy have been suggested as robust features for seizure detection and were used here to detect seizure-related components. The autocorrelation function of the signal was estimated to characterize cyclical patterns in the signal, the peaks in the autocorrelation function were identified, and the standard deviation and mean of the peak distances were calculated. Like spectral entropy, the entropy of the autocorrelation function may provide information on the regularity of the oscillation in the signal. A random signal will have a single peak at time 0. A fixed frequency oscillation will provide narrow and equal distance peaks in the function, resulting in slightly higher entropy. Changing spike times, phase shifts, or multiple oscillation patterns will create numerous irregular peaks in the autocorrelation function, leading to higher entropy values. Finally, the width of the autocorrelation function, a measure related to critical slowing[8], was calculated as previously described by Meisel et al. 2020[8].

We performed a preliminary evaluation of the extracted features to assess several aspects. First, the ***success proportion of feature extraction***: In several components, reliable features could not be extracted due to excessive noise. We quantified their proportion and omitted these from further analysis. Moreover, several features could not be calculated due to a short seizure or an insufficiently identified peak (under the defined parameters). Thus, samples with more than ten missing feature values were excluded from analysis; in the case of less than ten, the missing values were replaced with 0. Second, the mean difference between groups (clear vs. unclear and between bifurcation types) was explored to ensure it was as expected, mainly in morphological features. In this analysis, the ISI change features were more robust and consistent than amplitude modulation. The signal amplitude observed in surface EEG may not only relate to the properties of the seizure-initiating neuronal population. Instead, it may reflect the size of the neuronal population recruited as the seizure propagates, as demonstrated in supporting Figure 1C. Thus, in the current work, we decided to focus our automated classification on the distinction between seizures **with or without ISI slowing,** which may provide more reliable dynamic information in surface EEG data.

### Appendix 5: Classification considerations

To develop an automated classification system, we must first identify the components containing clear seizure bifurcations and then classify the bifurcation types for each seizure. Approaching this task, we must consider three main challenges in the data—first, ***noisy seizure data*** with possible outliers. To deal with this challenge, we scaled each feature column using RobustScalar (scikit-learn[9]), which is robust to outliers. Second, a ***relatively small sample size*** is prone to produce overfitting and reduces the generalization potential. To improve this, we first used both onset and offset samples to double the sample size. We added a categorical feature marking the location of the seizure to account for possible differences. Second, we used a random forest model[9] and limited the maximal tree depth to three, which has been demonstrated to be relatively resilient to overfitting[10]. ***Class imbalance***: both problems suffer from class imbalance. Simply training a classifier on such data will result in a classifier biased towards the majority class and having low sensitivity in the minority class. There are multiple ways to deal with this imbalance when building a classier[11]. In this work, we used the most straightforward approach: we set the class weights equal to the inverse proportion of the classes[12]. ***For model evaluation,*** we used a leave-one-out cross-validation approach which has been suggested as the proper evaluation in such cases[13]. Balanced accuracy was recorded for all patients[14]. In patients with both classes, the area under the ROC curve was calculated. The curve was not defined if only one class exited, making this calculation impossible. The distribution of all measures is reported. Lastly, we used a SHAP[15] analysis for each classifier to evaluate the contribution of each feature to each classification task.

### Supporting tables:

#### Supporting table 1: Onset selected components agreement

| | SN/SubH | SNIC | SupH |
|---|---|---|---|
| SN/SubH | **984** | 15 | 10 |
| SNIC | 3 | **54** | 0 |
| SupH | 1 | 0 | **65** |

#### Supporting table 2: Offset selected components agreement

| | FLC | SH/SNIC | SupH |
|---|---|---|---|
| FLC | **635** | 34 | 2 |
| SH/SNIC | 24 | **245** | 0 |
| SupH | 2 | 2 | **4** |

### Supporting table 3: Onset statistics

| Class | Parameter | MAP ratio | P-MAP | pd | ROPE |
|---|---|---|---|---|---|
| **SN/SubH** | Age 10-25 & 25-67 | 1.01 | 0.84 | 0.51 | 0.00 |
| | FBTCS & FIAS | 1.03 | 0.48 | 0.83 | 0.01 |
| | FBTCS & FAS | 1.00 | 0.99 | 0.62 | 0.03 |
| | FBTCS & UC | 1.05 | 0.46 | 0.87 | 0.01 |
| | FAS & FIAS | 1.02 | 0.69 | 0.75 | 0.01 |
| | FAS & FAS | 1.05 | 0.58 | 0.81 | 0.01 |
| | UC & FIAS | 0.98 | 0.77 | 0.71 | 0.00 |
| | Cortical dysplasia & Hippocampal sclerosis | 0.97 | 0.69 | 0.75 | 0.01 |
| | Female & Male | 1.00 | 0.99 | 0.53 | 0.00 |
| | Frontal & Temporal | 0.99 | 0.94 | 0.73 | 0.03 |
| | Length 60 < & > 60 seconds | 0.99 | 0.84 | 0.87 | 0.01 |
| | Bilateral & Left | 1.02 | 0.83 | 0.71 | 0.00 |
| | Bilateral & Right | 1.00 | 0.96 | 0.77 | 0.02 |
| | Right & Left | 1.03 | 0.33 | 0.96 | 0.00 |
| | Sleep stage I & Awake | 1.04 | 0.30 | 0.92 | 0.00 |
| | Sleep stage I & Sleep stage II | 1.00 | 0.98 | 0.55 | 0.01 |
| | Sleep stage II & Awake | 1.04 | 0.22 | 0.95 | 0.00 |
| | Sleep stage III/IV & Awake | 1.03 | 0.46 | 0.86 | 0.00 |
| | Sleep stage III/IV & Sleep stage I | 1.00 | 1.00 | 0.52 | 0.01 |
| | Sleep stage III/IV & Sleep stage II | 1.00 | 0.94 | 0.53 | 0.01 |
| **SNIC** | Age 10-25 & 25-67 | 1.25 | 0.83 | 0.79 | 0.13 |
| | FBTCS & FIAS | 0.00 | 0.98 | 0.59 | 0.22 |
| | FBTCS & FAS | 0.00 | 1.00 | 0.59 | 0.21 |
| | FBTCS & UC | 0.00 | 1.00 | 0.54 | 0.19 |
| | FAS & FIAS | 0.00 | 1.00 | 0.56 | 0.17 |
| | FAS & FAS | 0.00 | 1.00 | 0.56 | 0.17 |
| | UC & FIAS | 0.00 | 0.84 | 0.51 | 0.24 |
| | Cortical dysplasia & Hippocampal sclerosis | 0.00 | 0.95 | 0.82 | 0.30 |
| | Female & Male | 0.58 | 0.74 | 0.60 | 0.43 |
| | Frontal & Temporal | 0.00 | 0.99 | 0.66 | 0.32 |
| | Length 60 < & > 60 seconds | 0.04 | 1.00 | 0.81 | 0.11 |
| | Bilateral & Left | 0.67 | 0.76 | 0.63 | 0.38 |
| | Bilateral & Right | 0.00 | 1.00 | 0.73 | 0.16 |
| | Right & Left | 0.05 | 0.09 | 0.90 | 0.50 |
| | Sleep stage I & Awake | 0.00 | 0.07 | 0.90 | 0.19 |
| | Sleep stage I & Sleep stage II | 0.00 | 1.00 | 0.69 | 0.14 |
| | Sleep stage II & Awake | 0.00 | 0.53 | 0.84 | 0.36 |
| | **Sleep stage III/IV & Awake** | **0.00** | **0.00** | **1.00** | **0.01** |
| | Sleep stage III/IV & Sleep stage I | 0.00 | 1.00 | 0.94 | 0.01 |
| | Sleep stage III/IV & Sleep stage II | 0.00 | 1.00 | 0.98 | 0.01 |

| Class | Parameter | MAP ratio | P-MAP | pd | ROPE |
|---|---|---|---|---|---|
| SupH | Age 10-25 & 25-67 | 0.60 | 0.19 | 0.96 | 0.83 |
| | **FBTCS & FIAS** | **0.00** | **0.00** | **1.00** | **0.00** |
| | FBTCS & FAS | 0.00 | 1.00 | 0.91 | 0.02 |
| | **FBTCS & UC** | **0.00** | **0.00** | **1.00** | **0.00** |
| | FAS & FIAS | 0.00 | 0.00 | 0.99 | 0.08 |
| | FAS & FAS | 0.00 | 0.00 | 1.00 | 0.05 |
| | UC & FIAS | 1.30 | 0.74 | 0.87 | 0.07 |
| | Cortical dysplasia & Hippocampal sclerosis | 0.36 | 0.99 | 0.99 | 0.00 |
| | Female & Male | 0.93 | 0.99 | 0.56 | 0.30 |
| | Frontal & Temporal | 1.70 | 0.83 | 0.90 | 0.07 |
| | Length 60 < & > 60 seconds | 0.95 | 0.98 | 0.70 | 0.15 |
| | Bilateral & Left | 0.49 | 0.48 | 0.75 | 0.57 |
| | Bilateral & Right | 1.58 | 0.83 | 0.65 | 0.24 |
| | Right & Left | 0.39 | 0.16 | 0.92 | 0.76 |
| | Sleep stage I & Awake | 0.00 | 0.02 | 0.89 | 0.34 |
| | Sleep stage I & Sleep stage II | 0.00 | 1.00 | 0.56 | 0.16 |
| | Sleep stage II & Awake | 0.00 | 0.00 | 0.99 | 0.33 |
| | Sleep stage III/IV & Awake | 0.00 | 0.10 | 0.65 | 0.26 |
| | Sleep stage III/IV & Sleep stage I | 0.00 | 1.00 | 0.65 | 0.13 |
| | Sleep stage III/IV & Sleep stage II | 0.00 | 1.00 | 0.71 | 0.10 |

### Supporting table 4: Offset statistics

| Class | Parameter | MAP ratio | P-MAP | pd | ROPE |
|---|---|---|---|---|---|
| ISI Slowing | Age 10-25 & 25-67 | 1.19 | 0.52 | 0.89 | 0.01 |
| | FBTCS & FIAS | 0.96 | 1.00 | 0.51 | 0.31 |
| | FBTCS & FAS | 0.98 | 0.99 | 0.56 | 0.30 |
| | FBTCS & UC | 0.91 | 0.96 | 0.50 | 0.34 |
| | FAS & FIAS | 0.91 | 0.94 | 0.58 | 0.36 |
| | FAS & FAS | 0.77 | 0.81 | 0.58 | 0.41 |
| | UC & FIAS | 1.01 | 1.00 | 0.53 | 0.28 |
| | Cortical dysplasia & Hippocampal sclerosis | 0.82 | 0.76 | 0.64 | 0.44 |
| | Female & Male | 0.67 | 0.24 | 0.94 | 0.77 |
| | Frontal & Temporal | 0.50 | 0.31 | 0.88 | 0.74 |
| | Length 60 < & > 60 seconds | 1.14 | 0.54 | 0.90 | 0.00 |
| | Bilateral & Left | 1.45 | 0.49 | 0.93 | 0.02 |
| | Bilateral & Right | 1.71 | 0.22 | 0.98 | 0.00 |
| | Right & Left | 0.72 | 0.58 | 0.80 | 0.58 |
| | Sleep stage I & Awake | 0.01 | 0.31 | 0.87 | 0.55 |
| | Sleep stage I & Sleep stage II | 0.04 | 0.48 | 0.75 | 0.47 |
| | Sleep stage II & Awake | 0.85 | 0.81 | 0.74 | 0.48 |
| | Sleep stage III/IV & Awake | 0.44 | 0.70 | 0.71 | 0.47 |
| | Sleep stage III/IV & Sleep stage I | 0.00 | 1.00 | 0.62 | 0.24 |
| | Sleep stage III/IV & Sleep stage II | 0.51 | 0.79 | 0.59 | 0.39 |

Supporting table 5: Dynamotype rank and prevalence

| **Dynamotype** | **Total** | **Minimal codimension** | **Bifurcations required** |
|---|---|---|---|
| **SN/SubH – FLC** | 241 | 2 | 2 |
| **SN/SubH – SNIC/SH** | 83 | 3 | 2 |
| **SupH – SNIC/SH** | 6 | 3 | 3 |
| **SN/SubH – SupH** | 1 | 3 | 3 |
| **SupH – SupH** | 0 | 3 | 4 |
| **SNIC – FLC** | 9 | 4? | 3 |
| **SupH – FLC** | 14 | 4? | 5 |
| **SNIC – SNIC/SH** | 8 | 4? | 5 |
| **SNIC – SupH** | 0 | ? | ? |

### Supporting figures

#### S1: The labeling information

The following panels of information were presented to the rater:

(A) The EEG time series with an average reference montage

(B) The independent component's time series allowed us to identify the bifurcation morphology in the relevant component; component values are presented in procedure-defined units (PDUs).

(C) For additional component information used for classification, the reviewer was provided with additional component properties, including the onset and offset times evaluated, the spatial activation map of the components (left panel), the temporal activation map showing the signal power over time (right panel), and the power spectrum for each independent component (middle panel). The last component properties were presented using a costume code combined with EEGLAB visualization functions.

This figure demonstrates two components displaying prominent oscillatory seizure behavior. In IC2, we see an amplitude increase, whereas IC3 has a fixed amplitude. Observing the topographic activation of the components, we can see that the activation in IC2 is broader compared to IC3; hence, the amplitude increase may represent seizure propagation, rather than the pure dynamics of the epileptogenic zone.

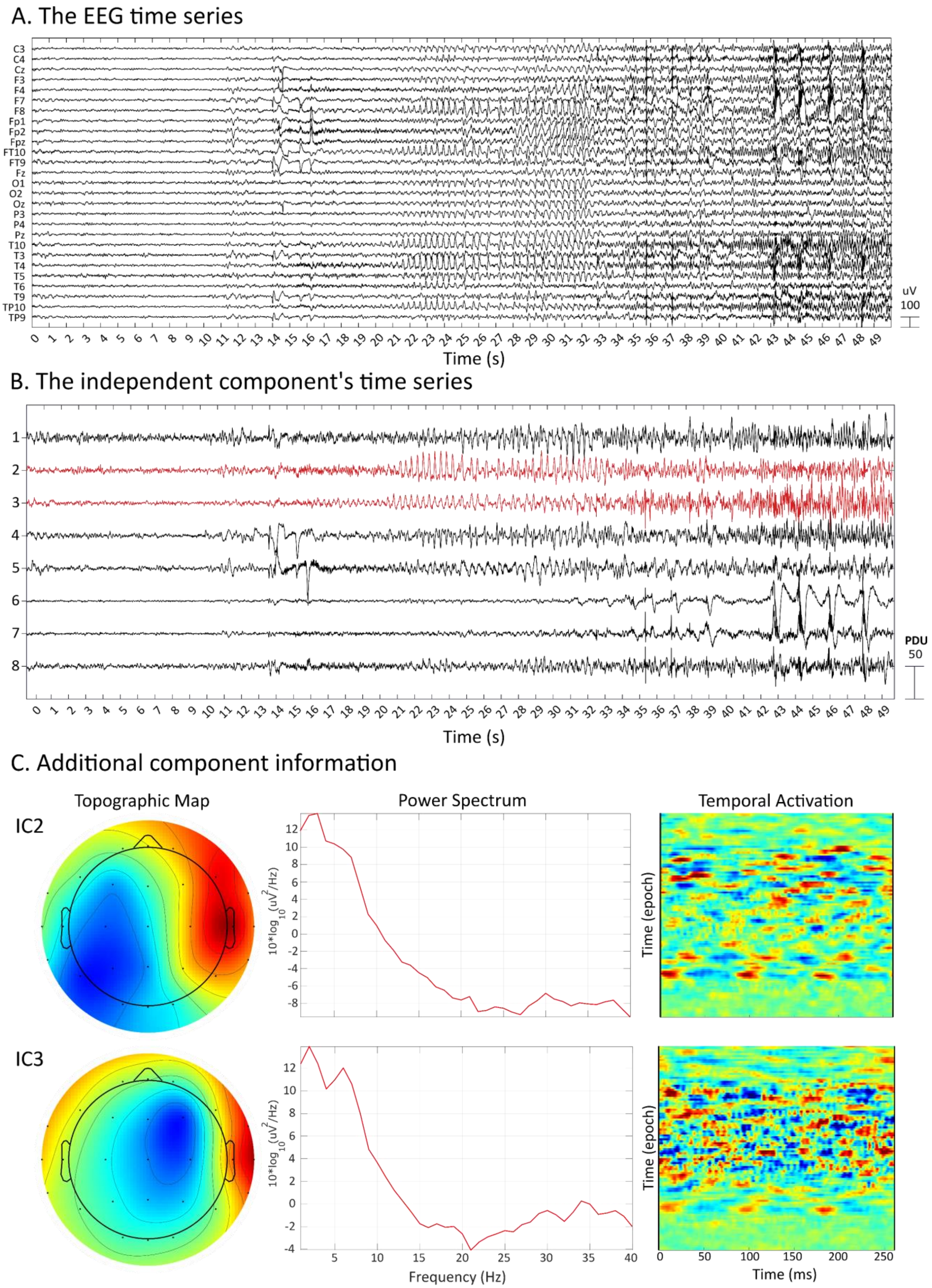
A. The EEG time series
C3
C4
Cz
F3
F4
F7
F8
Fp1
Fp2
Fpz
FT10
FT9
Fz
O1
O2
Oz
P3
P4
Pz
T10
T3
T4
T5
T6
T9
TP10
TP9
uV
100
Time (s)
B. The independent component's time series
PDU
50
Time (s)
C. Additional component information
Topographic Map
Power Spectrum
Temporal Activation
IC2
IC3
$10*\log_{10}(uV^2/Hz)$
Frequency (Hz)
Time (epoch)
Time (ms)

### S2: Onset examples

(A) All examples are presented in the procedure-defined units (PDUs). The result of the base transformation and the final scale depends on pre-whitening and the combination of the electrode weights. SN/SubH is the most prevalent onset type classified in our recordings. In some cases, the onset in this class is more subtle. The new labeling interface allowed us to better identify these changes in the better visually defined range.

(B) An example of an SN/SubH bifurcation with salient seizure onset activity: In the following figure, we can see the bifurcation time four seconds after the labeled onset time, within the allowed range. In some cases, this may be due to inaccuracies in the label marking the clinical onset of a seizure not being precisely timed with the electrographic start. In other cases, the labeled component may not correspond to the seizure onset zone, but may belong to a bifurcation in a propagation zone population.

(C) In some cases, amplitude increase will begin later, not directly at the onset. If the amplitude modulation is delayed from seizure onset, it will not be classified as SupH. In this example, the labeled bifurcation will be SN/SubH. We see the amplitude increase later in the seizure, perhaps due to a subsequent bifurcation or seizure propagation. Using the topographic activation map of this component (Figure 1C, IC2), we can see that this component has a relatively wide range of activation, which suggests that this may be due to the propagation of the seizure. In other cases (IC3), a more localized activation map will provide a better source for evaluating the bifurcation.

(D) The following seizure onset was defined as not clear. The neurologist described the onset pattern as "rhythmic theta waves." Slightly prior to five seconds, we can observe a baseline shift with reduced amplitude activity. Then, at 17 seconds, we can observe the beginning of an apparent oscillation in the theta range, which will fit the neurologist's description. In both cases, the bifurcation could be labeled as SN/SubH, but by our definition, these both occur outside of the labeling range, so this seizure is determined as unclear.

(E) The following demonstrates what we define as a brain component that occurs during seizure onset, but does not include a clear bifurcation.

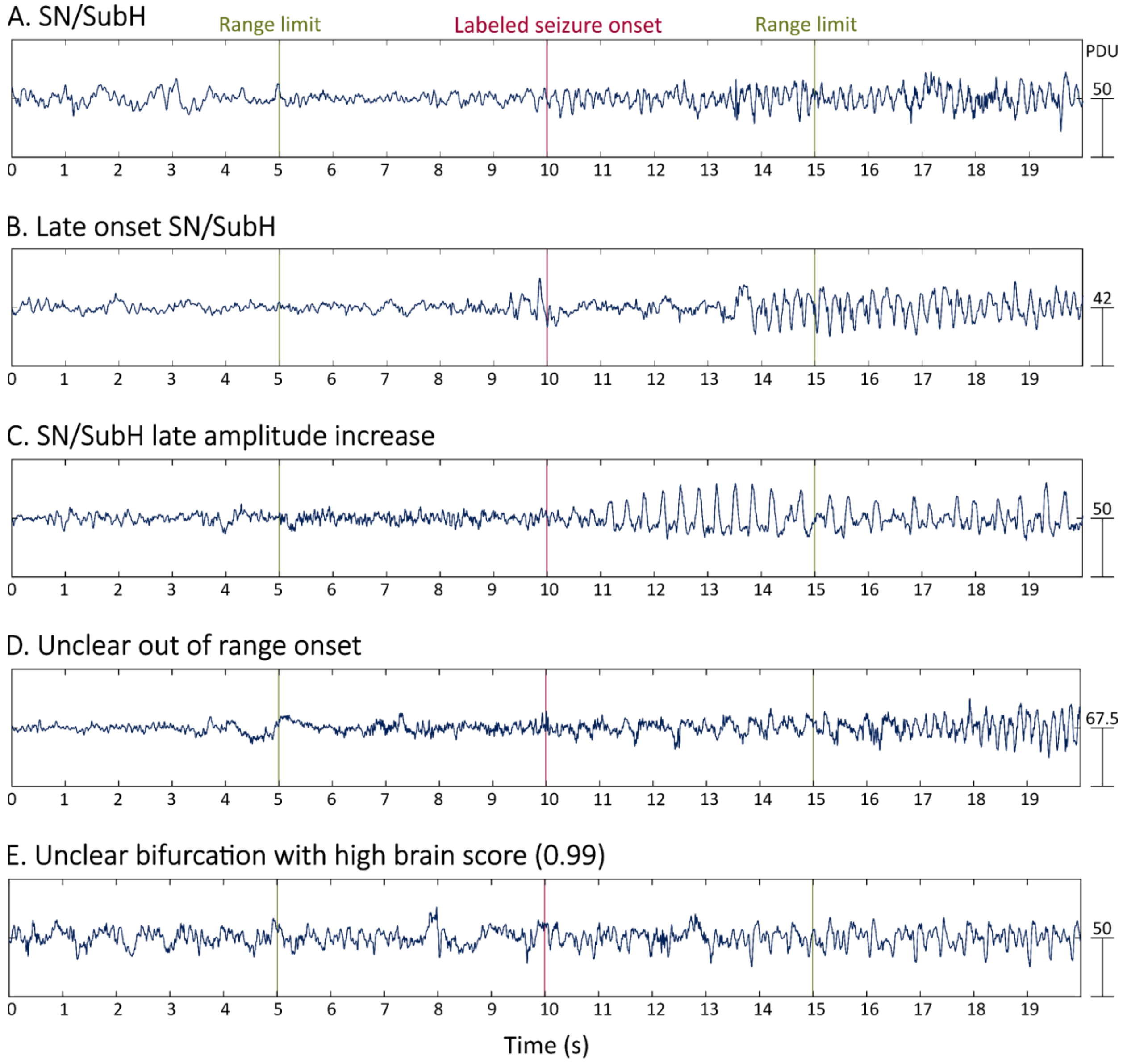

A. SN/SubH
Range limit
Labeled seizure onset
Range limit
PDU
50
B. Late onset SN/SubH
42
C. SN/SubH late amplitude increase
50
D. Unclear out of range onset
67.5
E. Unclear bifurcation with high brain score (0.99)
50
0 1 2 3 4 5 6 7 8 9 10 11 12 13 14 15 16 17 18 19
Time (s)


-

### S3: Offset examples

(A) We can see that the seizure ends at the end of the defined analysis run; this is visually classified, but will probably not yield a clear change point within the range of the automated, rule-based classification.

(B) In a similar matter as in the previous example, here, the seizure ends at the early edge of the analysis limit, and is not expected to yield a clear change point within the analysis range, even though this is a clear SH/SNIC bifurcation.

(C) The following demonstrates what we define as a brain component during seizure offset, yet it is hard to identify the exact termination time of the seizure oscillatory activity. Hence, we define it as an unclear bifurcation.

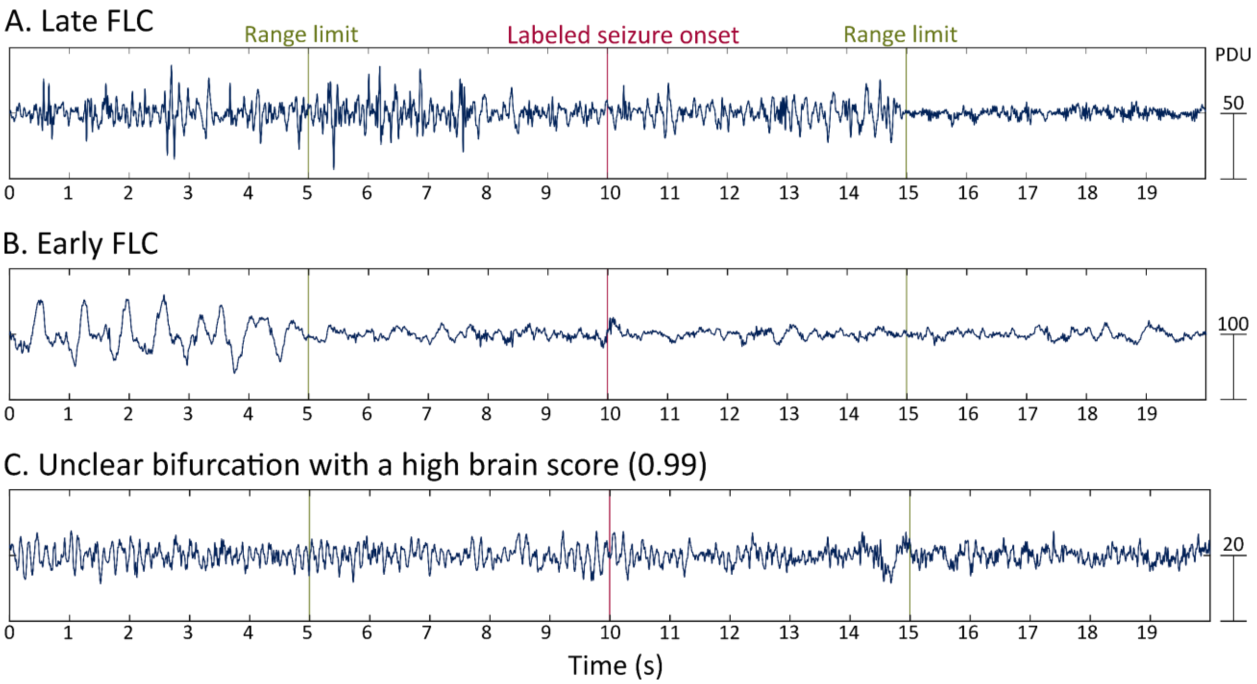

### S4: Score analysis and component selection

(A) The brain scores are presented overall and by the component number. The component number is related to the variability explained by the component, where lower values correspond to components explaining a more significant portion of the variability of the signal. Here, we can see that the overall selected component's score was lower for unselected components. This distinction was better preserved in components explaining a relatively large portion of the signal's variability. Additionally, the proportion of components with an apparent onset/offset was inversely related to their number. These suggest that producing additional independent components during a seizure may not give an additive value to the analysis.

(B) We used the earliest detected change point and the highest brain score in multiple labeled component cases. The decision flow used is illustrated in this panel.

A. Score analysis by bifurcation and component number

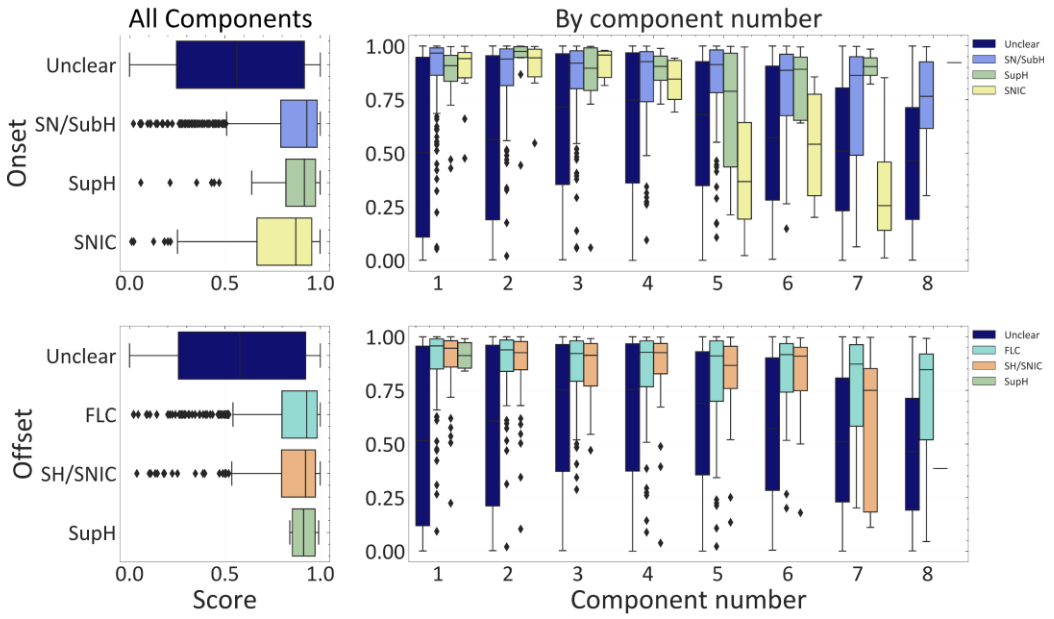


B. Component selection flow

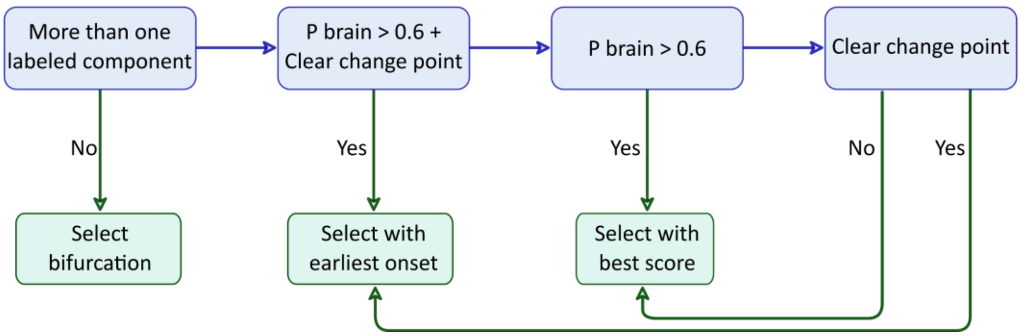

### S5: Detectability by onset and classification

The proportion of seizures with at least one component, including a clear bifurcation

(A) We can see that seizures originating from a posterior location have a higher probability of having a clear bifurcation both for onset and offset.

(B) We can see that seizures clinically classified as more extensive have a higher proportion of detectable bifurcations.

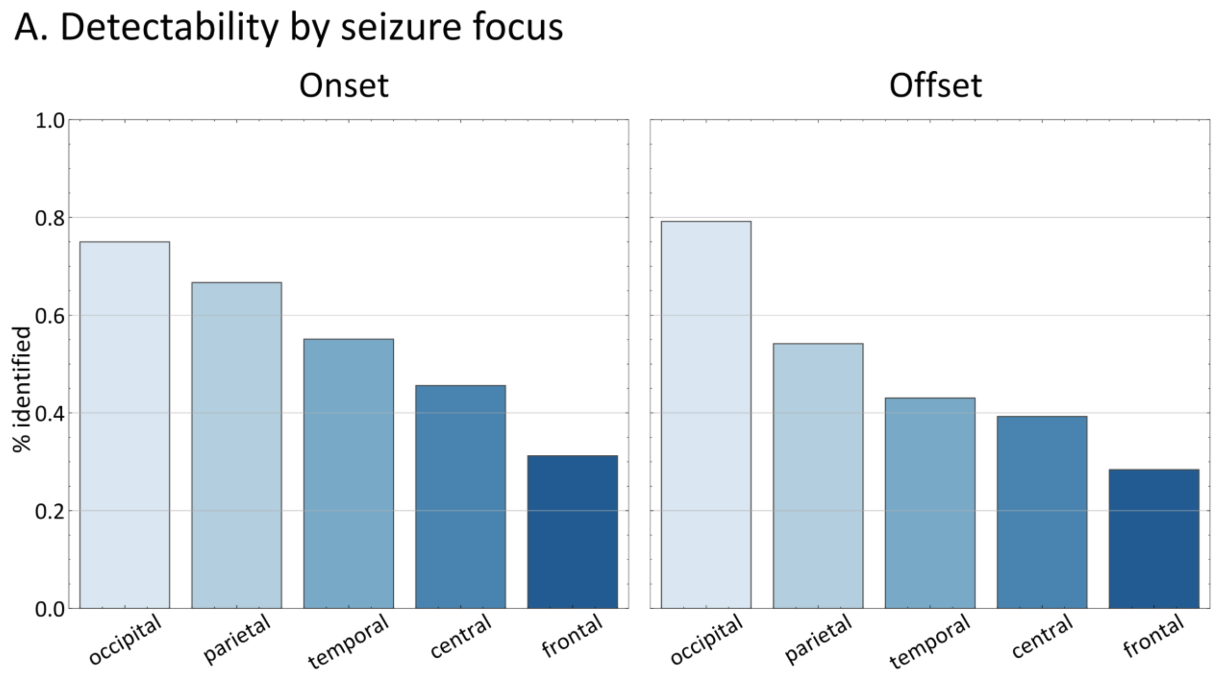


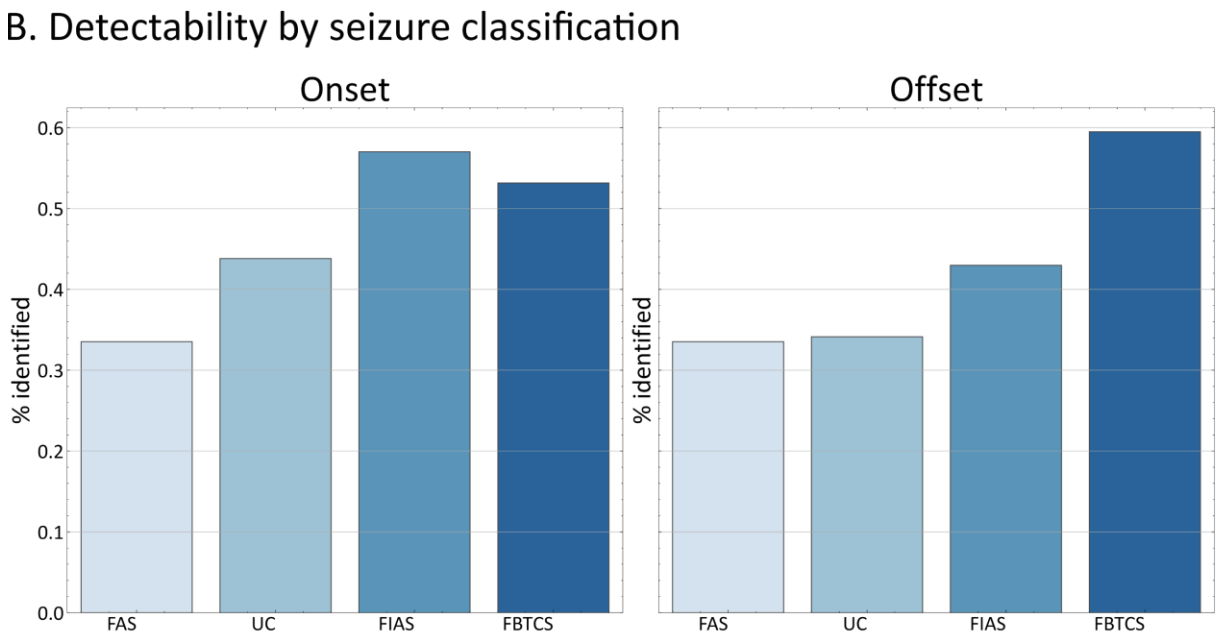
